# Neogene climatic fluctuations and poor connectivity with the centres of diversity shaped the Western Palearctic net-winged beetle fauna

**DOI:** 10.1101/2022.09.08.507108

**Authors:** Michal Motyka, Dominik Kusy, Renata Bilkova, Ladislav Bocak

## Abstract

Only twenty-two net-winged beetle species (Elateroidea: Lycidae) are known from the Western Palearctic region (WPR), i.e., less than 0.5% of the global lycid diversity and much fewer than from the similar latitudes of East Asia or Northern America. We use the comprehensive distribution data and the molecular phylogeny of ∼400 world lycids, including fourteen European species, to provide a new perspective for understanding the structure and evolution of this group in the WPR. All Mediterranean species represent deeply rooted lineages with relatives in Eastern Asia. These species occur in relictual ranges close to the family’s Pleistocene refugial edge. The phylogeny points to the loss of biological connection with East Asia since the Mid Miocene. A third of WPR species is widespread in Central and Northern Europe, reaching Eastern Asia, some of them possibly younger elements of the European fauna. Unlike relatively high diversity in the Eocene amber, the extant net-winged beetles represent a small fraction of elateroid diversity in the WPR and are generally rare. Therefore, we assume that most WPR species are relics trapped in Mediterranean refugia since the onset of the Plio-Pleistocene cooling and are critically endangered by the ongoing loss of suitable habitats.

## INTRODUCTION

Europe and Northern Africa have been severely affected by Pliocene and Pleistocene climatic fluctuations (Mossbrugger *et al*., 2005; Hughes & Gibbard, 2018; Cheddadi *et al*., 2021). Their beetle fauna (∼30,000 spp.) is only slightly more diverse than the Northern American fauna (25,500 spp., but many are still undescribed). It is also much smaller if compared with the even less known Eastern Asian fauna (∼50,000 spp.; Löbl & Smetana 2007; Miller 2002). Although biotic responses to past shifts in climate have been intensively studied, most analyses concentrated on closely related species or populations with a time frame of fewer than 1 million years (my; Medail & Diadema, 2009; Hampe & Jump, 2011). Less information is available on diversity changes over long periods (e.g., Harzhauser *et al*., 2016) and the phylogenetic relationships of European and Eastern Asian beetle faunas (e.g., Sanmartín *et al*. 2001; Sota *et al*., 2008, 2022; Brunke *et al*., 2019).

Here, we use net-winged beetles (Lycidae) as a model for inferring the evolutionary history of beetles in the Western Palearctic region (WPR). This elateroid family contains approximately 4,500 species worldwide, but only twenty-two in the WPR (Tabs 1, S1; Bocak & Bocakova, 1987; Bocakova & Bocak, 2007; Kazantsev, 2011; Masek *et al*., 2018). We assume that the ranges of net-winged beetles are primarily defined by at least periodical accessibility of moist organic material exploited by larvae with mouthparts adapted to liquid food (Bocak & Matsuda, 2003). Like larvae, the desiccation-prone soft-bodied adults also prefer moist places, and only some species tolerate semiarid conditions (some Calochromini, Lycini, and Metriorrhynchini; Masek *et al*., 2018). The family is highly diverse in the tropics, where only a fraction of the diversity has been described (Motyka *et al*., 2021a). The Holarctic net-winged beetle fauna is relatively poor in species number (80 Nearctic and ∼300 Palearctic species; Miller, 2002; Bocakova & Bocak, 2007). Nevertheless, it contains most subfamilies and tribes (Kusy *et al*., 2019). In contrast to the rather thoroughly inventoried WPR fauna, more species will undoubtedly be described from East Asia. T. Nakane and other Japanese entomologists have studied the net-winged beetles in eastern Asia since the 1950s (e.g., Nakane, 1969, Matsuda, 2010). Recently, numerous species have been described in China (e.g., Kazantsev, 1997, 2004; Li *et al*., 2015, 2017).

Recent intensive investigations of the Eocene amber fauna have expanded information on the structure of the European fauna 55.8–33.9 million years ago (Alexeev, 2017; Tab. S2). The early European Cenozoic is characterized by a gradual transition from the Lower Eocene climatic optimum, 52–55 million years ago (Mya), to a striking climate change at the Eocene/Oligocene boundary (33.7 Mya) and gradual cooling in later periods (Zachos *et al*., 1996, 2001; Szwedo & Sontag, 2009; Szawaryn, 2021). Subfamilies and tribes of net-winged beetles supposedly originated long before the Eocene, and one subfamily was recorded in Burmese amber (Motyka *et al*., 2017, 2021b; Bocak *et al*., 2019; Tihelka *et al*., 2019). The Eocene net-winged beetles were already reported in the early 20^th^ century (Klebs, 1910; Kleine, 1940). Kazantsev (2013) reviewed the Eocene amber fauna, corrected some misidentifications, and identified four lycid species in three genera (Supplementary Table S2). Kazantsev (2019) and Kazantsev & Perkovsky (2022, Tab. S2) described the other two species making five lycid genera and six species, of which three genera are extinct. The richness of the Eocene European fauna may be related to the climatic optimum with substantially higher temperature and high precipitation, i.e., the conditions favourable for net-winged beetles (Masek *et al*., 2018). Although the dating remains uncertain, the Baltic amber fossils indeed predate the closure of the Turgai gap and capture the fauna thriving before the Oligocene cooling (Zachos *et al*., 1996, 2001; Brikiatis, 2014; Brunke *et al*., 2019).

The present study builds on the previous molecular phylogenies of Lycidae. Kusy *et al*. (2019) redefined subfamilies and tribes using phylogenomic data, and Motyka *et al*. (2021b) built the phylogeny of >500 species, i.e., about 12% of world fauna utilizing the combination of the phylogenomic backbone and Sanger data. The DNA data on Chinese fauna were reported by Li *et al*. (2015, 2017), Motyka *et al*. (2017), and Kusy *et al*. (2021). We add further sequences for the Western Palearctic species. We assume that the net-winged beetles have been sufficiently sampled to estimate the relationships of WPR species in the context of the whole Palearctic region. We specifically try to explain present-day biogeographical patterns in Europe, North Africa, the Caucasus, and northern Iran using DNA-based relationships within the worldwide net-winged beetle fauna. We investigate the evolution of our model group in the context of European paleoclimate and the earlier defined refugia of the diversity in the Western Palearctic region.

## MATERIAL AND METHODS

### Study area

We study the fauna of the Western part of the Palearctic region (WPR) following the Palearctic region’s borders defined earlier by Löbl & Smetana (2007) and Udvardy (1975). Alternatively, our study area covers the western parts of the Saharo-Arabian (only NW Africa) and Palearctic regions (Holt *et al*., 2013). The study area is delimited in the east by the Ural Mountains and the wide strip of deserts in central Asia (Kazakhstan, Uzbekistan, and Turkmenistan), Iran, Gulf states, and the north of the Arab Peninsula, where no net-winged beetles occur (Fig. 1). The northern part of the defined eastern border is permeable (West Siberian taiga and Kazakh forest-steppe).

**Figure 1.**
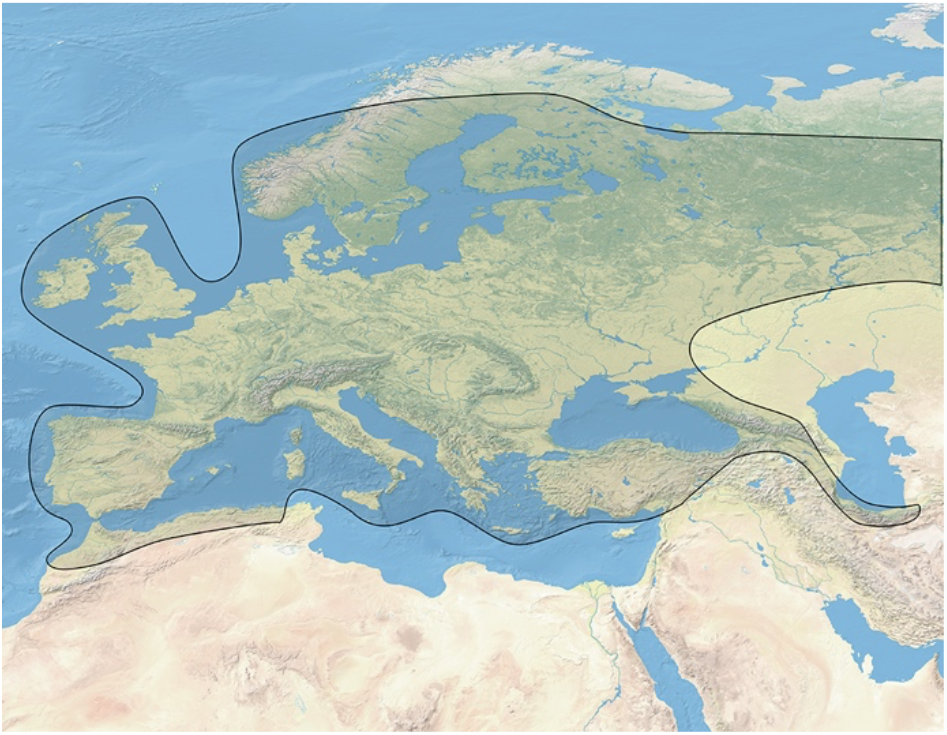
Delimitation of the Western Palearctic Region (WPR) and distribution of net-winged beetles.

The southern part of the WPR includes several ecoregions with generally threatened fauna: Mediterranean conifer and mixed forests (North-western Africa), Southwest Iberian Mediterranean sclerophyllous and mixed forests, Pyrenees conifer and mixed forests, Cantabrian mixed forests (Spain, Portugal); South Apennine mixed montane forests (South Italy); Anatolian conifer and mixed deciduous forests (Southern Turkey); Caucasus mixed forests, Crimean Submediterranean forest complex, Euxine–Colchic deciduous forests (Caucasus and adjacent regions); Caspian Hyrcanian mixed forests (Azerbaijan, Iran; Olson *et al*., 2001).

### Material

Fourteen of twenty-two WPR species were sequenced for three mitochondrial fragments. The list of the WPR species in the analysis is given in Tab. S1, along with voucher codes that designate terminals in phylogenetic trees. There are a few missing species, mostly rare species, and some are known only from type specimens described more than 100 years ago. (Tab. 1). The distribution data were assembled from major European museum collections (see depositories), the Fauna Europea/GBIF database, and the inaturalist.org webpage.

**Table 1.**
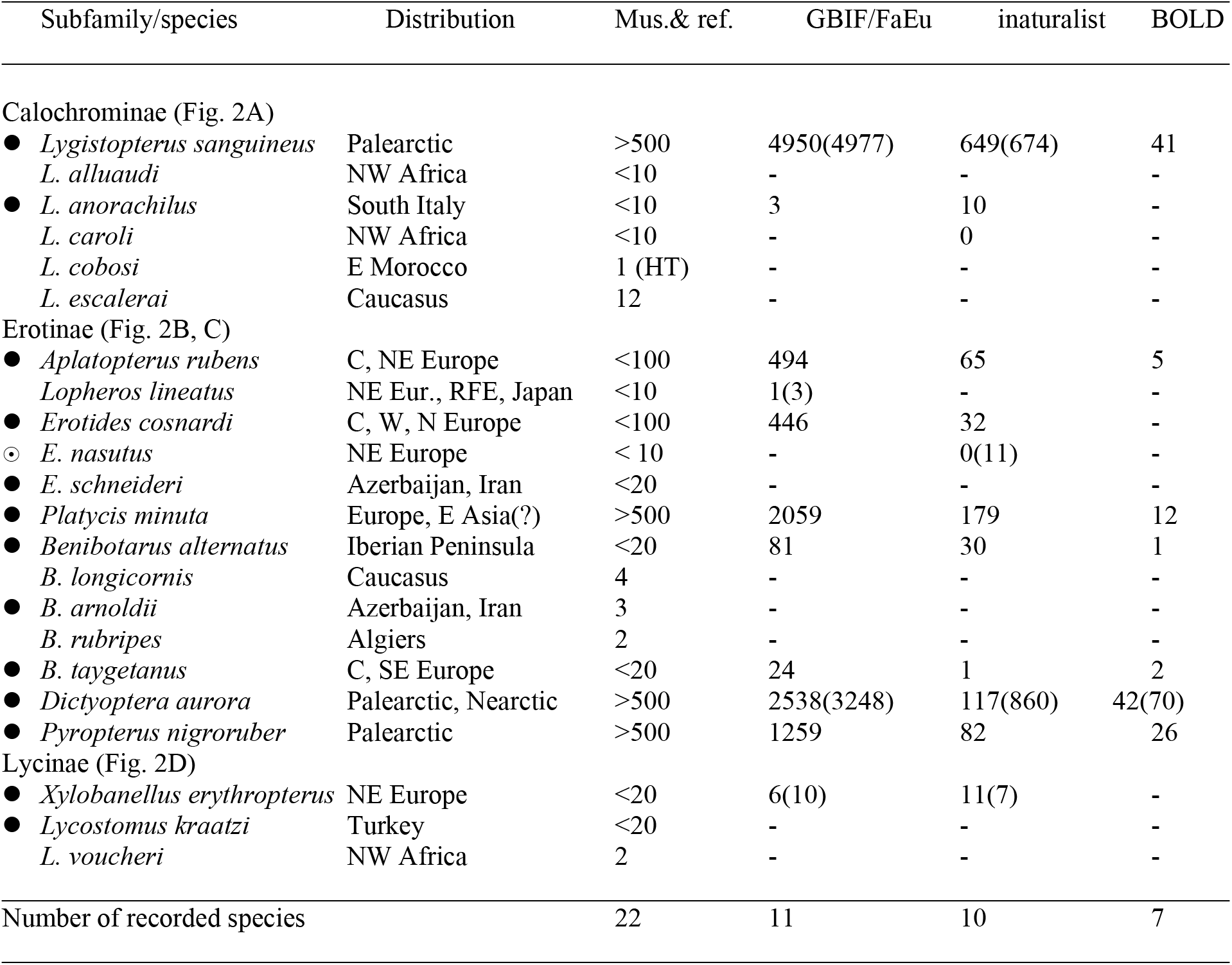
The list of European species, distribution data, and estimation of their relative abundance measured by the presence in studied European collections (see Methods), Fauna Europea/GBIF database (www.gbif.org), inaturalist (www.inaturalist.org), and the BOLD records (www.ibol.org). The numbers report georeferenced records in the GBIF database (de Yong 2016), all observations reported by inaturalist, and the number of sequences deposited in the Barcode of Life database. All data as of March 4^th^, 2022. Numbers refer to western Palearctic records, all records incl. non-European are given in parentheses. The classification overview and detailed distribution data are shown in Supplementary Table S1. Legend: • Western Palearctic species included in the analysis; ⊙ conspecific individuals of non-Western Palearctic origins included in the study; - stands for species not recorded by the database; HT holotype; RFE Russian Far East.

### Depositories

National Museum, Prague, Czech Republic, Moravian Museum, Brno, Czech Republic, Museum and Institute of Zoology, Warszawa, Poland; National Natural History Museum, Paris, France; Hungarian Natural History Museum, Budapest, Hungary; Zoological Museum at the Humboldt University, Berlin, Germany; State Natural History Museum, Stuttgart, Germany; Museum of Natural History, London, UK, Natural History Museum, Basel, Switzerland, senior author’s collection.

### Laboratory procedures

Molecular data, phylogenetic analyses, and zoogeography The phylogenetic investigation is based on earlier published *cox1, nadh5*, and *rrnL* mtDNA, SSU, and d2 loop of LSU rRNA sequences. The data for most WPR species were explicitly produced for this study. The laboratory methods were described by Bocak *et al*. (2008).

We assembled a matrix of 432 lycid terminals and *Iberobaenia minuta* Bocak *et al*., 2016, a sister to the Lycidae, as an outgroup (Bocak *et al*., 2016; Tab. S1). The length invariable protein-coding *cox1* and *nadh5* sequences were aligned using TRANSALIGN (Bininda-Emonds, 2005) and the rRNA fragments using default parameters of MAFFT v.7.2 (Katoh & Standley, 2013).The individual gene alignments were concatenated into a supermatrix (Tab. S3). The substitution models were identified using ModelFinder (Kalyaanamoorthy *et al*., 2017). We estimated maximum likelihood (ML) trees using IQ-Tree v.1.8.1 (Nguyen *et al*., 2015) and the option -g, allowing us to constrain the deep topology by relationships inferred by the earlier phylogenomic analyses (Kusy *et al*., 2019). The bootstrap support (BS) was obtained using UFboot (Hoang *et al*., 2018) with 5,000 replicates.

We calibrated the tree by the earlier inferred age of the family (Bocak *et al*., 2016; McKenna *et al*., 2019). The position of two fossil records (*Pseudaplatopterus* Kleine, 1940 as a sister of the *Aplatopterus* Reitter, 1911 + *Lopheros* LeConte, 1881 clade and *Helcophorus* Fairmaire, 1891 as a sister to *B. alternatus* (Fairmaire, 1856) was used for the verification of the dating analysis.

The Beast v.1.8 analysis (Drummond *et al*., 2012) of the unlinked mtDNA dataset was set to the HKY+I+G model, and the birth-death speciation prior; genes and codon positions were partitioned in the (1+2),3 scheme. The four independent analyses were run for 5*10^7^ generations with a sampling frequency of 10,000 generations. All independent runs were checked for ESS values and plateau phases using TRACER v.1.681 (Rambaut *et al*., 2018). LogCombiner v.1.8.1 was used to discard 33% of the trees as a burn-in and to produce final log files. The maximum credibility tree was generated with TREEANNOTATOR v.1.8.1 (Drummond *et al*., 2012). All trees were visualized using Figtree v.1.4.4 (https://github.com/rambaut/figtree) and Adobe Illustrator.

## RESULTS

The taxonomic overview, taxonomic authorities, and distribution of the Lycidae WPR fauna are given in detail in the Supplementary Text. Basic information is summarized in Tab. 1 and Figs 2A–D. We identified six widespread, cold-tolerant taxa with ranges reaching beyond 60ºN latitude [*L. sanguineus* (L., 1757), *P. nigroruber* (De Geer, 1774), *D. aurora* (Herbst, 1784), *P. minuta* (F., 1787), *E. cosnardi* (Chevrolat, 1831), and *A. rubens* (Gyllenhal, 1817)] and three species tolerant to continental low winter temperatures, with the northern distribution limits between 52ºN and 60ºN but limited to north-eastern Europe [*X. erythropterus* (Baudi di Selve, 1871), *E. nasutus* (Kiesenwetter, 1874), *L. lineatus* (Gorham, 1883); Fig. 2C, D]. Six of these species are distributed to the eastern Palearctic; three are known only in the WPR (*A. rubens, E. cosnardi*, and *P. minuta*). One species is distributed in Central Europe north of the Alps and the Balkans [*B. taygetanus* (Pic, 1905); Fig. 2B]. It does not occur in areas with the continental climate and only marginally reaches the south (a single record from the Taygetos in the Peloponnese, the nearest northern localities known from Romania and Bosnia; Fig. 2B). All these species are considered widespread.

**Figure 2.**
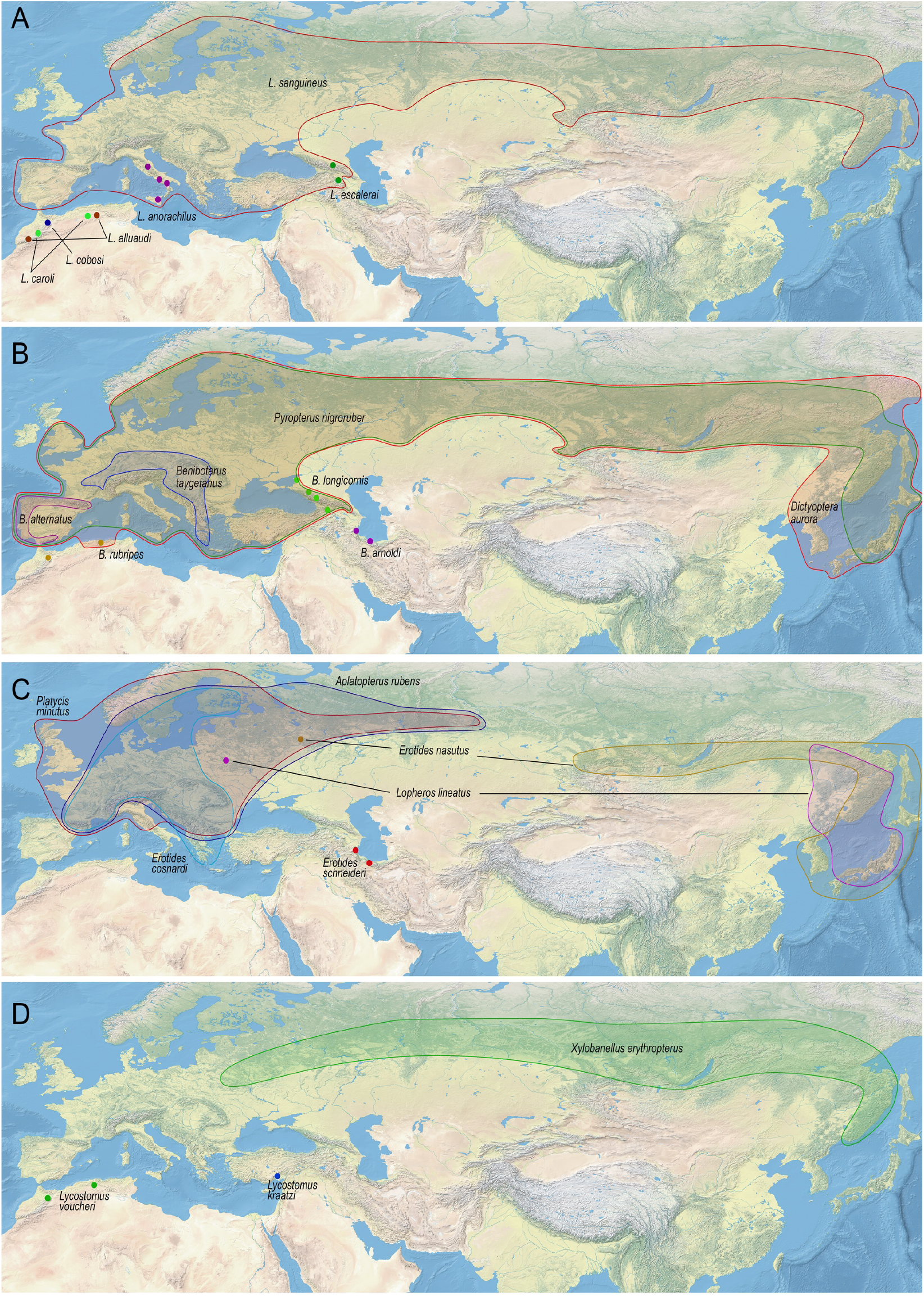
A – Distribution of Calochrominae in the Palearctic region (*Lygistopterus*). B – Erotinae: Dictyopterini in the Palearctic region (*Benibotarus, Dictyoptera*, and *Pyropterus*). C – Erotinae: Erotini in the Palearctic region (*Aplatopterus, Erotides, Lopheros*, and *Platycis*). D – Lycinae: Conderini (*Xylobanellus*) and Lycini (*Lycostomus*) in the Palearctic region.

Twelve species have a relictual distribution, several have only been recorded from very restricted ranges and in a single or few specimens. *Lygistopterus caroli* (Bourgeois, 1881), *L. cobosi* (Pardo Alcaide, 1961), *L. alluaudi* (de Peyerimhoff, 1925), *Lycostomus voucheri* (Bourgeois, 1905), and *Benibotarus rubripes* (Pic, 1897) occur in Mediterranean conifer and mixed forests; *B. alternatus* in Pyrenees conifer and mixed forests and Cantabrian mixed forests; *Lyg. anorachilus* Ragusa, 1883 in South Apennine mixed montane forests; *Lyg. escalerai* Pic, 1942, *B. longicornis* (Reiche, 1887) in Caucasus mixed forests, Crimean Submediterranean forest complex, and Euxine–Colchic deciduous forests; *B. arnoldii* (Barovsky, 1932), *E. schneideri* (Kiesenwetter, 1874) in Caspian Hyrcanian mixed forests; *Lyc. kraatzi* Bourgeois, 1882 in Anatolian conifer and mixed deciduous forests (Tab. 1, Fig. 2A–D).

The ranges of some genera are disjunctive (*Benibotarus* Kôno, 1932, *Lycostomus* Motschulsky, 1861). Similarly, *L. lineatus* and *E. nasutus* occur in two widely separated populations (Euclidean distance ∼6,000 km and 3,000 km, respectively, Fig. 2B). North-eastern European and North Asian temperate to subarctic forests ensure the connectivity between the WPR and East Asia (6 spp., Fig. 2A–D), and the arid areas of South-western Asia are impermeable for net-winged beetles.

### Phylogenetic relationships to worldwide net-winged beetles

The WPR fauna substantially differs from the World in the subfamily and tribal structure. The most diverse net-winged beetle lineages (Metriorrhynchinae: Metriorrhynchini, >1400 spp.; Platerodini, >800 spp. worldwide) do not occur in the WPR. Erotinae: Erotini, 54 spp. worldwide, and Dictyopterini, 73 spp. worldwide, represent a substantial part of net-winged beetle diversity in the WPR (altogether 13 spp., 59% of species versus 3% worldwide). Similarly, Calochrominae is proportionally well represented (6 spp. in the WPR, 27% of diversity compared with 7% representation worldwide). The remaining three species belong to Lycinae: Conderini (1 sp.) and Lycini (2 spp.).

The WPR fauna shows relationships to Eastern Asia. We have not found any connection between the Afrotropical and Western Palearctic fauna. Several species from the Nearctic regions have sister species in Eastern Asia and generally represent more shallow splits than the WPR species (Figs 3, S1). *Platycis minuta, D. aurora, A. rubens, L. kraatzi, P. nigroruber*, and *B. alternatus* (the last two as serial splits), *B. arnoldii* + *B. taygetanus*, and the clade *E. nasutus* (*E. schneideri, E. cosnardi*) are sisters to more extensive East Asian monophyla (Figs 3, S1). Only *Xylobanellus erythropterus* is a terminal within a dominantly East Asian clade. The ancestor with the WPR distribution can be hypothesized for two species pairs (*E. cosnardi, E schneideri*, and *B. taygetanus, B. arnoldii*).

**Figure 3.**
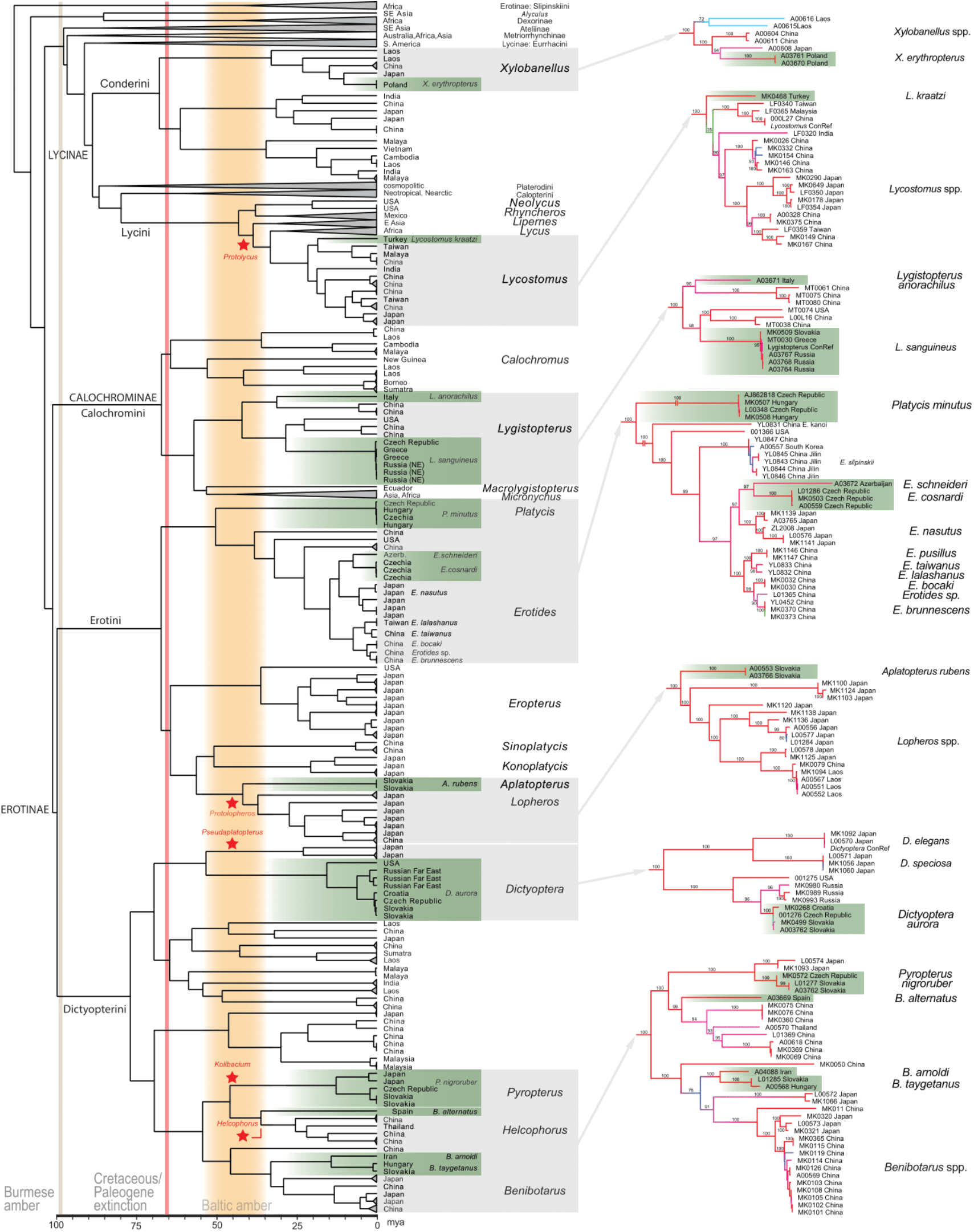
Left – Dated tree of the family Lycidae, bars show 95% confidence intervals, the letters a–p designate splits listed in Tab. 2; stars designate the position of Eocene amber records listed in Tab. S2. Right – Focal topologies with the Western Palearctic representatives; phylograms with UFboot values above branches; the complete trees in Figs S1, S2. The subtrees are connected to corresponding parts of the dated tree by arrows.

The dating analysis is validated by Baltic amber lycids records (Figs 3A, S2), and the splits between WPR net-winged beetles and their closest relatives are summarized in Tab. 2. The results show the absence of speciation in the WPR since the Upper Miocene, i.e., the split between *E. schneideri* and *E. cosnardi* from the Hyrcanian Caspian Forest and central Europe, respectively. The youngest species splits were recovered between Japanese and European species of *Xylobanellus* Kleine, 1930 (10.4 may) and between two species occurring in the southern part of the WPR region that diverged in the Mid to Late Miocene (14.39 and 7.45 Mya, Tab. 2). Most species stand alone as deeply rooted lineages separated from their closest relatives 19 to 51 Mya (Tab. 2).

**Table 2.** The mean age and 95% confidence intervals for splits of the Western Palearctic species and their sister lineages in a million years. The letters designate splits shown in Figure 3.

| Species | Geographic origins of sister taxa |  | Mean time | 95% confidence intervals |  |
| --- | --- | --- | --- | --- | --- |
| a/ *Xylobanellus erythropterus* | Poland | Japan | 10.40 | 4.38 | 16.27 |
| b/ *Lycostomus kraatzi* | Turkey | China | 18.61 | 12.70 | 24.58 |
| c/ *Lygistopterus anorachilus* | Italy | China | 31.58 | 15.25 | 46.43 |
| d/ *L. sanguineus* | Europe | China | 28.73 | 17.91 | 39.88 |
| e/ *Platycis minuta* | Europe | China & USA | 50.96 | 39.87 | 62.77 |
| f/ *Erotides cosnardi* | Europe | Azerbaijan | 7.45 | 3.90 | 11.62 |
| g/ *E. nasutus* | Japan | Europe/Azerbaijan | 11.93 | 8.42 | 15.41 |
| h/ *Erotides* spp. | W Palearctic | China, Japan | 14.80 | 11.06 | 18.68 |
| i/ *Aplatopterus rubens* | Europe | Japan, Laos | 42.30 | 32.87 | 52.51 |
| j/ *Dictyoptera aurora* | Holarctic | Japan | 53.93 | 39.62 | 67.16 |
| k/ *D. aurora* (intraspecific) | Palearctic | Nearctic | 15.75 | 26.06 | 7.18 |
| l/ *D. aurora* (intraspecific) | Europe | Russian Far East | 6.23 | 9.70 | 3.27 |
| m/ *Pyropterus nigroruber* | Europe | Europe, China | 46.37 | 23.19 | 49.79 |
| n/ *P. nigroruber* (intraspecific) | Europe | Japan | 12.84 | 5.09 | 22.01 |
| o/ *Benibotarus alternatus* | Spain | China | 36.59 | 23.19 | 49.79 |
| p/ *B. taygetanus* | Europe | Azerbaijan | 14.39 | 4.53 | 27.79 |

The phylogenetic analyses indicate genetic divergence between Asian, Nearctic, and WPR populations of *D. aurora* (Fig. 3) and between East Asian and WPR populations of *P. nigroruber* (Fig. 3, S1, S2). Conversely, we did not recover genetic differentiation within European populations of *L. sanguineus*, which has a similar range as the species mentioned above and for which we sequenced populations separated by >2,000 km (Tab. S1, Fig. S1).

## DISCUSSION

### Shared distribution patterns of net-winged beetles

The late Cenozoic and Quaternary aridization and cooling (Mossbrugger *et al*., 2005; Hughes & Gibbard, 2018) drove the extinction of the warm and humid climate fauna and flora and the WPR was more affected than Eastern Asia due to different geomorphological conditions (Svenning, 2003; Burke *et al*., 2018; Chen et al. 2018). However, little is known about the different responses of various groups. For example, soldier beetles, well-represented in Baltic amber, still dominate the extant elateroid WPR’s fauna (Löbl & Smetana, 2007; Alexeev, 2017). Conversely, net-winged beetles are generally rare and resemble in the extent of faunal turnover rove beetles (Tab. 1; Bocak & Bocakova, 1987; Brunke *et al*., 2019). More than half of WPR species have a relictual distribution (Fig. 2A–D). Therefore, we assume that they were severely affected by the Pliocene-Pleistocene climatic changes. The lycid larvae depend on liquid food and have a limited dispersal capacity (Bocak & Matsuda, 2003; Li *et al*., 2015; Jiruskova *et al*., 2019; Motyka *et al*., 2021a, b). As a result, most species cannot readily exploit ex-situ refugia.

Based on the geographic distribution, we can identify four species groups in the WPR fauna of Lycidae. The widely distributed, euryvalent, cold-tolerant species occur in temperate to subarctic deciduous and conifer forests. The ranges of these species cover large areas north of the Alps. Presumably, they were able to react to the climate-driven loss of suitable habitats by shifting their ranges during cold periods. Only these species colonized the Northern European forests established after the latest retreat of the European continental glacier (Hughes & Gibbard, 2018). Here, we assign six species (27% of diversity): *L. sanguineus, D. aurora, A. rubens, Pl. minuta, E. cosnardi*, and *Pyr. nigroruber*. These six species also occur, although more rarely, in Southern Europe, supposedly serving as a refugium during the last glacial maximum (Fig. 2A– D).

Over half of the lycids in the WPR seems to be trapped in small refugia located within the less suitable semiarid regions along the retreating southern edge of the family range (12 spp., 55% of the WPR diversity; Fig. 2A–D): Northwest African mountains (5 spp.), Caspian Hyrcanian forests (2 spp.), the Caucasus (2 spp.), the Pyrenees, the west and south-west of the Iberian Peninsula (1 sp.), southern Italy (1 sp.) and southern Turkey (1 sp.). We found the complete species turnover between these areas and it confirms their status as distinct ecoregions and their function as refugia during climatic fluctuations (Olson *et al*., 2001, Keppel et al. 2012). Most of these species are very rare; some Moroccan and Algerian species have not been collected for the past hundred years – *Lycostomus voucheri* and *Lygistopterus cobosi*. Others are known in a few individuals from recent field research (*Lyg. caroli, Lyg. alluaudi, Benibotarus rubripes*). Similarly, the species from the Caucasus and Caspian Hyrcanian forests are rare (Tab. 1).

Although we could intuitively expect that at least some of these species were separated and diversified due to the fragmentation of ranges (Cardoso & Vogler, 2005; Gomez-Zurita *et al*., 2000), the occurrence of the common ancestor in the WPR is indicated only for two pairs of species: *B. taygetanus* / *B. arnoldii* and *E. cosnardi* / *E. schneideri* (Figs 3, S1, S2). These species pairs are distributed along the putative dispersal route between the WPR and Eastern Palearctic regions. The two sequenced species of *Lygistopterus* are very distant, and each has its closest relative in the Eastern Palearctic region. North African *L. caroli* and *L. alluaudi* are morphologically divergent and were placed in a separate genus (Motyka *et al*., 2017). Similarly, *B. longicornis, B. alternatus*, and *B. rubripes* have their close relatives only in the Eastern Palearctic region. The analysed *Lycostomus* represents a deeply rooted lineage and does not have close relatives in the WPR (Fig. 3).

The third distinct group contains species occurring from north-eastern Europe to north-east Asia, i.e., in the regions with dominantly continental, cold-winter climates and sufficiently high precipitations keeping organic substrates wet for most of the year. Three species, 14% of WPR’s fauna, are assigned to this group: *L. lineatus, E. nasutus*, and *X. erythropterus*. All are rare in the WPR and have only been recorded from a few localities in north-western Poland (Bialowieza), Lithuania, Belarus, and Central Russia (the Moscow region; Kazantsev 2011). The European parts of their ranges were mainly under the ice sheet during the last glacial maximum. They probably followed the retreating glacier with forest habitats southeast of the adjacent tundra (Hewitt, 2000; Hughes & Gibbard, 2018). The available records for *L. lineatus* and *E. nasutus* indicate highly fragmented ranges of these species, although we cannot exclude that the observed pattern is a sampling artifact (Fig. 2C; Kazantsev, 2011).

One species does not fit either group. *B. taygetanus* occurs in central Europe and the Balkans. It is not a widespread species, and although distributed partly in the area very close to the limits of Quaternary glaciation limits (Hughes & Gibbard, 2018), it did not colonize northern Europe, unlike the first group. *B. taygetanus* is rare and only marginally reaches western Europe and the Mediterranean. Its sister species, *B. arnoldii*, is known for the Caspian Hyrcanian Forest (Figs 2B, 3, S1, S2), and their closest relatives are known from Japan and China.

The aridization of the Mediterranean and the Caspian region possibly affected the viability and persistence of local net-winged beetle species. Although the loss of suitable conditions is rarely complete, and some patches with favourable microclimate (ravines, windward slopes) are usually preserved, the net-winged beetles in the southern part of the range are rare, and some are known only in a few specimens (Tab. 1). It is well supported by the molecular topology (Fig. 3) and morphology-based placement that their relatives occur not closer than in the Himalayas (*Helcophorus* spp., *L. caroli* group), the Hindukush (*Lycostomus*), and Southwestern China (Sichuan, Yunnan; *Pyropterus, Helcophorus-Pyropterus*; Kazantsev, 2004, 2011; Bocakova & Bocak, 2007; Masek *et al*., 2018). The rareness of all species, the fragmentation of their ranges, and geographical disconnection from their relatives show that North African and South European lycids are highly endangered by the loss of habitats due to aridization and, recently, the human-driven degradation of oak and cedar forests to shrub-like communities. The species cannot effectively react to the loss of habitats due to the absence of corridors, low dispersal propensity, and the distance of potential ex-situ refugia (Ashcroft, 2010).

### Age and origins of WPR lycids

The WPR fauna represents a subset of the Holarctic lycid diversity with a long history since at least the mid-Eocene, as documented by five genera from three tribes preserved in Baltic amber (Supplementary Table S2; Miller, 2002; Motyka *et al*., 2021b; Kazantsev 2013). Alexeev (2017) reported 614 beetle species in Eocene amber, of them four Lycidae, and further two lycids were described later (Tab. S2; Kazantsev, 2019; Kazantsev & Perkovsky, 2022). Lycidae represented 0.5% to 1% of the Eocene beetle fauna. As all lycid adults are poorly flying and primarily inactive, we assume their overrepresentation in amber is improbable.

We assume that net-winged beetles represent a remnant of the earlier rich Tertiary fauna, and we base our assumption on the following arguments. The extant WPR net-winged beetle fauna is species-poor, and most species are rare (∼0.6‰ of the WPR beetle fauna (Löbl & Smetana 2007), 0.1‰ of beetle observations in the GBIF database; de Jong, 2016). It is a much sparser occurrence than in Baltic amber (0.5–1% of inclusions; Alexeev, 2017; Tab. S2). Notably, the diversity of lycids in Baltic amber is comparable with the extant Japanese fauna of the warm and humid climate resembling those of Europe in the Eocene (Nakane, 1969; Zachos *et al*., 1996, 2001; Burke *et al*., 2018). Secondly, some genera present in Eocene amber went extinct, and others are represented in the WPR by one or two extant species (Tabs S1, S2; Bocakova & Bocak, 2007; Kazantsev, 2013). Finally, over half of WPR species occur in very restricted ranges and are separated by intervening unfavourable conditions from the nearest regions with the occurrence of their relatives (Fig. 2A–D).

The WPR fauna is zoogeographically isolated, and we did not identify any group of Afrotropical origin in the Mediterranean, Caucasus, and northern Iran. In the Mid to Late Miocene, the net-winged beetles missed the Afro-Arabian-Mediterranean dispersal corridors connecting Africa, Arabia, and Eurasia that were used by some vertebrates (Tejero-Cicuendez, 2022; Jolivet & Facenna, 2000). Although the Sahara Desert was temporarily covered by shrub vegetation in the Pliocene and Pleistocene (Cheddadi *et al*., 2021), the Sub-Saharan species did not reach the WPR.

The fauna of Eastern North America (mainly the Appalachians) shares with Europe the three genera, *Lopheros* Le Conte, 1881, *Erotides* Waterhouse, 1879, and *Dictyoptera* Latreille, 1829, and two, *Lopheros* and *Erotides*, are absent in the Western USA (Miller, 2002), possibly due to aridization of the region as was proposed for plants (Donoghue & Smith 2004). In contrast with the absence in the Western USA and diversified fauna of the erotines in Baltic amber, we do not indicate that the De Geer and Thulean land bridges played any role in the colonization of the Nearctic region by Erotinae, eventually Calochrominae (Fig. 3; Enghoff 1995). The Erotinae fauna of Eastern North America might have been established by dispersal through the Bering route available during the Paleogene, ∼58 Ma (Brikiatis, 2014) and later (Sota *et al*., 2008, 2022). The analysed Nearctic species are sisters to East Asian lycids and split from them later than the WPR relatives (Fig. 3).

The WPR fauna is characterized by the dominance of Erotinae (14 spp. of ∼160 spp. worldwide), one of the three earliest lineages of the family (Kusy *et al*., 2019) that is most diverse in the temperate Holarctic and uncommon in the tropics (Kazantsev, 2004; Li *et al*., 2017; Masek *et al*., 2018). Other groups, the Calochrominae (6 spp.) and Lycinae: Conderini, are also deeply rooted. (Fig. 3, Motyka *et al*., 2017, 2021b; Masek *et al*., 2018). Calochrominae is cosmopolitan, with its origin in Southeast Asia (Motyka *et al*., 2017), and Conderini is mainly Oriental, with a few species in the Palearctic region. The Lycini (2 spp., Figs 2D, 3) has been identified as a Neotropic group dispersing to Asia via the Beringia route and, later, to Africa, eventually western Asia (Kusy *et al*., 2021). Several highly diversified tribes are absent in the WPR, and they belong to groups of presumed Neotropical/Mesoamerican (Lycinae: Platerodini) or Australian (Metriorrhynchini: Cautirina) origins (Bocakova 2001; Sklenarova *et al*., 2013; Masek *et al*., 2018). Possibly these lineages did not occur in the Eastern Palearctic when dispersal routes to the west were effectively open.

The European fauna is close to temperate and subtropical eastern Asia, belonging to the same zoogeographical region. Still, we do not find evidence for a long-term connection between these subregions. The Turgai Sea and the southern fragmented island chain limited the dispersal between Asia and Europe during most Paleogene (Sanmartín *et al*., 2001; Brikiatis, 2014). The continental sea dried only in the Lower Oligocene. The newly opened route between the WPR and East Asia was also used by mammals and ground beetles (Hooker & Dashzeveg, 2003; Sota *et al*., 2022). Later, the continual northern forest belt might have contributed to the dispersal of some cold-resistant species (*L. lineatus, E. nasutus, X. erythropterus*) that split from relatives in the Upper Miocene or later (Fig. 3). We found a pretty ancient split between genetically distinct European and Japanese populations of *P. nigroruber* and *D. aurora* (estimated splits 12.8 and 15.8 Mya, respectively). In these cases, the Siberian dispersal route is highly probable (Fig. 2B– D). The northern and southern routes connect the WPR region to the distinct Sino-Japanese and Sino-Himalayan faunas defined based on the analysis of plant distribution (Chen *et al*., 2018) but not so apparent in the East Palearctic net-winged beetle fauna.

The Balkanatolia might contribute to the dispersal of some Asian taxa to Europe through a southern route shortly before the Grande Coupure at the Eocene-Oligocene boundary (Licht *et al*., 2022). According to the dating analysis (Figs 3A, S2), Erotinae was already in Europe in the Mid to Upper Eocene, as documented by Baltic amber. The Baltic amber *Protolycus gedaniensis* Kazantsev, 2019 (Lycini; Kazantsev, 2019) is a member of older fauna than the extant *Lycostomus kraatzi* included in the analysis. We assume that the Mediterranean endemics or their ancestor species established the range in WPR shortly before the origin of Baltic amber fauna (Fig. 3, Motyka *et al*., 2021b). Later, the low connectivity between East Asia and the WPR might have been supported by the aridization of western and southern Asia (Retallack, 2001). *Lygistopterus* Mulsant, 1838 colonized the WPR in the Early Oligocene (28.7, 31.6 Mya, Tab. 2). The origin of *Helcophorus*, recently reported from Rovno amber (Kazantsev & Perkovsky, 2022), was recovered slightly younger than the fossil record (Fig. 3A). At least since the Miocene, south-western and central Asia represent a wide dry barrier separating the extant Sino-Himalayan and Western Palearctic faunas. We have not found any species with a recent split from their East Asia relatives that could disperse through these semi-desert areas (Figs 1, 3).

## CONCLUSION

Speciation capacity, extinction rates, and dispersal propensity characterize the potential of a species for long-term survival. Many beetles reacted to the changing Pliocene and Pleistocene climates with diversification (Gomez-Zurita *et al*., 2000; Cardoso & Vogler, 2005). We found a different pattern in net-winged beetles. Most European species are deeply rooted, isolated lineages, and only in two cases did we find species pairs with a putative European ancestor. Almost all European species remain in the relict ranges, with distant relatives far from their extant range. Considering the structure of the Eocene Baltic amber and extant Eastern Nearctic faunas, we assume that the WPR lycids represent the last surviving species of an original Tertiary Western Palearctic fauna. Their actual status was affected by their biological characteristics and the Pliocene to Pleistocene climatic changes that similarly devastated the tree Cenozoic diversity of Western Palearctic forests (Svenning, 2003).

## ACKNOWLEDGMENTS

This study would not have been possible without the help of numerous colleagues who provided us with several rare European species: S. Fabrizzi, L. Dembicky, J. M. Gutowski, and A. Ocampo. The field research in China was partly conducted by Y. Li, V. Kubecek, and K. Sklenarova and was generously supported by H. Pang (Guangzhou); the Japanese Society for the Promotion of Science supported the collecting of the senior author in Japan.

## CONTRIBUTION STATEMENT

MM: conceptualization, formal analysis, investigation, writing – review & editing. DK: conceptualization, formal analysis, investigation, writing – review & editing. RB: Investigation, writing – review & editing. LB: conceptualization, investigation, writing – original draft, visualization, funding acquisition. M.M. and D.K. contributed equally to this work.

## FUNDING

The Czech Science Foundation project 22-33714S supported this work.

## DATA AVAILABILITY

The analyzed dataset was deposited in the Mendeley data repository (doi:10.17632/b67237fxb5.1).

## CONFLICT OF INTERESTS

We have no competing interests.

## SUPPORTING INFORMATION

**Supplementary text**. Taxonomic overview and distribution of the Lycidae fauna of the Western Palearctic region.

**Table S1**. The list and distribution of the extant West Palearctic Lycidae with the geographic origins of the samples in the analysis.

**Table S2**. The updated list of fossil Lycidae.

**Table S3**. The partition scheme, the best substitution models and information about IQ-TREE run.

**Figure S1**. Molecular phylogenetic reconstruction of Lycidae relationships using the maximum-likelihood approach.

**Figure S2**. Time-calibrated, maximum clade credibility tree computed using BEAST.

## Notes

### Competing Interest Statement

The authors have declared no competing interest.

