## SUPPORTING INFORMATION for "Neogene climatic fluctuations and poor connectivity with the centres of diversity shaped the Western Palearctic net-winged beetle fauna"

### Supplementary information

**Supplementary text**
Taxonomic overview and distribution of the Lycidae fauna of Western Palearctic region.

**Supplementary tables**
**Table S1.** The list and distribution of the extant West Palearctic Lycidae with the geographic origins of the samples in the analysis.
**Table S2.** The updated list of fossil Lycidae.
**Table S3.** The partition scheme, the best substitution models and information about IQ-tree run.

**Supplementary figures**
**Figure S1.** Molecular phylogenetic reconstruction of Lycidae relationships using maximum-likelihood.
**Figure S2.** Time-calibrated, maximum clade credibility tree computed using BEAST.

**Suplementary text**
***Taxonomic overview and distribution of the Lycidae fauna of Western Palearctic region***

Calochrominae: Calochromini
*Lygistopterus* Mulsant, 1838
All WPR Calochrominae are placed in *Lygistopterus*; two of six extant representatives, *L. sanguineus* and *L. anorachilus*, were analyzed (Tab. 1). Despite the phenotypic similarity, *L. sanguineus* and *L. anorachilus*, are distantly related. *L. cobosi* Pardo Alcaide 1961 and *L. escalerai* are phenotypically similar to the sequenced species (Cobos 1961, Terzani et al. 2015) and possibly have belonged to a widespread mimetic ring as similarly colored species do not occur in the eastern part of the Palearctic region. Only *L. sanguineus* is widespread, and others occur in small ranges close to the southern limits of the tribal range and are all represented by a few specimens in collections (Fig. 2, Tab. 1).

Erotinae: Erotini
Six Erotini species have been recorded from the WPR, five were included in the analyses (Tab. 1). We found high diversity in Western, Central, Northern, and Northeastern Europe (5 spp., two of them, *Platycis* and *Aplatopterus,* representing monotypic genera, other placed in *Erotides* – 2 spp. and *Lopheros* – 1 sp.). Only some of these species marginally reach the Mediterranean in southern France, northern Italy, and the Balkans (Fig. 3). Two species, both extremely rare in Europe, have disjunctive ranges. *Lopheros lineatus* was reported from Bialowieza Puszta in Poland and the Russian Far East, over 6,000 km away. The closest population of *Erotides nasutus*, recently collected in the Moscow region, is known in Altai, over 3,000 km further to the east (Fig. 3). *E. schneideri* is known from several localities in the Caspian Hyrcanian Forest and is closely related to central European *E. cosnardi*. All three Erotides form a clade (Fig. 6).

Erotinae: Dictyopterini
Seven Dictyopterini species are known from the WPR, and five of them were analyzed. The tribe contains two widespread species, *Dictyoptera aurora* and *Pyropterus nigroruber*, both having a range >$10^7$ $km^2$. *D. aurora* is the only WPR species with the Holarctic distribution, but the populations from Northern America are genetically very distinct (max. uncorrected cox1-5' pairwise distance 12.8%; Fig. S2, Tab. S4) and may represent a separate species that should bear the name *D. coccinata* (Say, 1835). *Pyropterus* (1 sp. in the analysis) is a sister to East Asian and Himalayan *Helcophorus* (Fig. 6). Four species occur close to the southern rim of the tribal range: *B. alternatus* in Iberia, *P. nigroruber*, *P. rubripes* in Northern Africa, *B. longicornis* in the Caucasus, and *B. arnoldi* in the Caspian Hyrcanian Forest (Fig. 4). *B. alternatus* was recovered as a sister to *Pyropterus,* and we found that *B. arnoldi* is a sister to *B. taygetanus* (Fig. 6).

Lycinae: Lycini
Two species of *Lycostomus* have been recorded from the WPR (Fig. 5); one of them, *L. kraatzi* was analyzed and found at the deep split within the genus (Fig. 6). Both species are relics of the Mediterranean region, *L. voucheri* is extremely rare and was unavailable for analysis.

Lycinae: Conderini
Only *Xylobanellus erythropterus* occurs in Northeastern Europe (Bialowieza Puszta, Latvia, Northern Russia, unconfirmed record from Finland), and its range reaches the Russian Far East (Fig. 5). The congeners occur in Japan, Southern China (undescribed species) and Southeast Asia.

### Supplementary tables

**Table S1.** The list and distribution of the extant West Palearctic Lycidae with the geographic origins of the samples in the analysis. The sequenced species are designated by black dots if they were available from the Western Palearctic Region. A circle with a small dot designates the species known from Europe but sequenced only from Japan.

Calochrominae Lacordaire, 1857
tribe Calochromini Lacordaire, 1857
Lygistopterus Mulsant, 1838
●*Lygistopterus sanguineus* (L., 1758) Europe, Turkey, East Kirghizstan, East Kazakhstan, West and East Siberia, Far East (Primorskij Kraj, Sakhalin). Samples: Czech Republic, Greece, NE Russia
*L. alluaudi* (Peyerimhoff, 1925) Morocco
●*L. anorachilus* Ragusa, 1883 Sicily, Southern Italy. Sample: Italy.
*L. caroli* (Bourgeois, 1882) Morocco, Algiers
*L. cobosi* Pardo Alcaide 1961 E Morocco
*L. escalerai* Pic, 1942 (Georgia, Ossetia, Armenia)

Erotinae LeConte, 1881
tribe Erotini LeConte, 1881
●*Aplatopterus rubens* (Gyllenhal, 1817). Europe, Southwestern Siberia. Samples: Slovakia.
●*Erotides (Glabroplatycis) cosnardi* (Chevrolat, 1831) Ukraine (Transcarpathia). Central and Western Europe, Southern Scandinavia. Samples: Czech Republic.
●*E. (Glabroplatycis) nasutus* (Kiesenwetter, 1874)
NE Europe (Moscow Oblast), Southwestern Siberia, Russian Far East, Japan, Korea. Samples: unavailable from the Western Palearctic Region, sampled only from Japan.
●*E. (Glabroplatycis) schneideri* (Kiesenwetter, 1878) Caspian Hyrcanian forests. Sample: Azerbaijan.
*Lopheros lineatus* (Gorham, 1883) Poland (Bialowieza), Russian Far East, Japan, northeastern China.
●*Platycis minutus* (Fabricius, 1787) Europe, West Siberia. Samples: Czech Republic, Hungary.

tribe Dictyopterini Kleine, 1928
*Benibotarus (s. str.) longicornis* (Reiche, 1878). Eastern Black Sea coast (Krasnodar Kraj), Abkhazia, (Sukhumi), Georgia (Batumi, Borjomi).

●*B. (s. str.) alternatus* (Fairmaire, 1856) Portugal, Spain, the Pyrenees. Sample: NW Spain.
●*B. (Sibetarus) arnoldii* (Barovskij, 1932) Caspian Hyrcanian forests. Sample: Iran.
●*B. (Sibetarus) taygetanus* (Pic, 1905) Ukraine (Transcarpathia). Central and southeastern Europe. Samples; Hungary, Slovakia.
*B. rubripes* Pic, 1897 Algiers (holotype only).
●*Dictyoptera aurora* (Herbst, 1784) Europe, West and East Siberia, Russian Far East, Eastern Kazakhstan, Japan, Korea, Iran, North Africa (Algeria), North America. Samples: Czech Republic, Croatia, Slovakia, Russian Far East, USA.
●Pyropterus nigroruber (Degeer, 1774) Europe, Siberia, Russian Far East (northern Sakhalin), Japan. Samples: Czech Republic, Slovakia, Japan.

subfamily Lycinae Laporte, 1838
tribe Conderini Bocak et Bocakova, 1990
●*Xylobanellus erythropterus* (Baudi di Selve, 1872) Poland, North Eastern Europe Urals, West and East Siberia Russian Far East. Samples: Poland.
tribe Lycini Laporte, 1838
●*Lycostomus kraatzi* Bourgeois, 1882 Turkey. Sample: Turkey.
*L. voucheri* Bourgeois, 1905 Algiers, Morocco.

**Table S2.** The updated list of fossil Lycidae. The Eocene amber species are designated by black dots.

**Lycidae Laporte, 1840 Lycinae Laporte, 1840**
**Dexorinae Bocak & Bocakova, 1989**
Burmolycini Bocak, Li et Ellenberger, 2019
*Burmolycus* Bocak, Li et Ellenberger, 2019: 152.
Type species *B. compactus* Bocak, Li et Ellenberger, 2019
*compactus* Bocak, Li et Ellenberger, 2019: 152. Cretaceous Burmese amber (97 mya).

*Murcybolus* Li, Tihelka Huang et Cai, 2021
Type species *M. longiantennus* Li, Tihelka Huang et Cai, 2021
*longiantennus* Li, Tihelka Huang et Cai, 2021. Cretaceous Burmese amber (97 mya).

Cretolycini Tihelka, Huang et Cai, 2019: 263
*Cretolycus* Tihelka, Huang et Cai, 2019: 263.
Type species C. compactus Tihelka, Huang et Cai, 2019

*praecursor* Tihelka, Huang et Cai, 2019: 264. Cretaceous Burmese amber (97 mya).

**Lycinae Laporte, 1840**
Lycini Laporte, 1840
*Protolycus* Kazantsev, 2019: 329.
Type species *Protolycus gedaniensis* Kazantsev, 2019
●*gedaniensis* Kazantsev, 2019: 330. Eocene Baltic amber (~35 mya).

Platerodini Kleine, 1929

*Plateros* Bourgeois, 1879: xix.

Type species *Eros brasiliensis* Lucas, 1857

*jardinesi* Kazantsev, 2020, Mexican amber (20 mya).

Leptolycini Leng et Mutchler, 1921

*Electropteron* Kazantsev, 2020

Type species *Electropteron avus* Kazantsev, 2020

*avus* Kazantsev, 2020 Miocene Dominican amber (~20 mya).

*Cessator* Kazantsev, 2009: 93.

Type species Cessator luquillonis Kazantsev, 2009.

*brodzinskyi* Ferreira et Ivie, 2017: 58. Miocene Dominican amber (~20 mya).

**Erotinae LeConte, 1881**

Erotini LeConte, 1881

= Lopherotini Kazantsev, 2012

*Pseudaplatopterus* (*Pseudaplatopterus*) Kleine, 1940: 179.

Type species *P. ascheelei* Kleine, 1940

●*ascheelei* Kleine, 1940: 179. Eocene Baltic amber.

*Protolopheros* Kazantsev, 2013: 94.

Type species *P. hoffeinsorum* Kazantsev, 2013.

●*hoffeinsorum* Kazantsev, 2013: 97. Eocene Baltic amber.

Dictyopterini Houlbert, 1922

*Helcophorus* Fairmaire, 1881: cxxix.

Type species *H. miniatus* Fairmaire, 1881.

= *Hiekeolycus* Winkler, 1987: 66. Type species *H. berendti* Winkler, 1987.

●*berendti* (Winkler, 1987): 67. Eocene Baltic amber.

●*ekaterinae* Kazantsev & Perkovsky, 2022: 85. Eocene Rovno amber.

*Kolibacium* Winkler, 1987: 62.

Type species *K. balticum* Winkler, 1987.

= *Pietrzeniukia* Winkler, 1987: 68. Type species *P. kunowi* Winkler, 1987

●*balticum* Winkler, 1987: 63. Eocene Baltic amber.

= *Pietrzeniukia kunowi* Winkler, 1987: 70.

**Taxa excluded from Lycidae**

*Prototrichalus* Molino-Olmedo, Ferreira, Branham et Ivie, 2020: 2; described in Metriorrhynchini Kleine, 1926, now Tenebrionoidea *incertae sedis* (Bocak et al. 2022).

*Prototrichalus meiyingae* Molino-Olmedo, Ferreira, Branham et Ivie, 2020: 6. Cretaceous Burmese amber.

*Prototrichalus milleri* Molino-Olmedo, Ferreira, Branham et Ivie, 2020: 6. Cretaceous Burmese amber.

*Prototrichalus sepronai* Molino-Olmedo, Ferreira, Branham et Ivie, 2020: 6. Cretaceous Burmese amber.

Table S3. The partition scheme, the best substitution models and information about IQ-tree run.

| Subset | Seqs | Sites | Infor | Invar | Model | Name |
|---|---|---|---|---|---|---|
| 01 | 365 | 656 | 376 | 211 | GTR+F+I+G4 | *rrnL* |
| 02 | 299 | 68 | 39 | 20 | GTR+F+I+G4 | *rrnL* tRNA |
| 03 | 190 | 1904 | 256 | 1519 | SYM+I+G4 | *18S* |
| 04 | 194 | 647 | 123 | 480 | SYM+I+G4 | *28S* |
| 05 | 344 | 1014 | 829 | 125 | GTR+F+I+G4 | *nadh5* |
| 06 | 332 | 292 | 195 | 61 | TPM3u+F+I+G4 | *nadh5* tRNAs |
| 07 | 339 | 780 | 491 | 249 | GTR+F+I+G4 | *cox1* |
| 08 | 248 | 257 | 201 | 41 | TIM+F+I+G4 | *cox2* |
| 09 | 255 | 65 | 37 | 22 | GTR+F+I+G4 | *cox1* tRNA |
| 10 | 254 | 123 | 92 | 17 | TIM+F+I+G4 | *nadh1* |

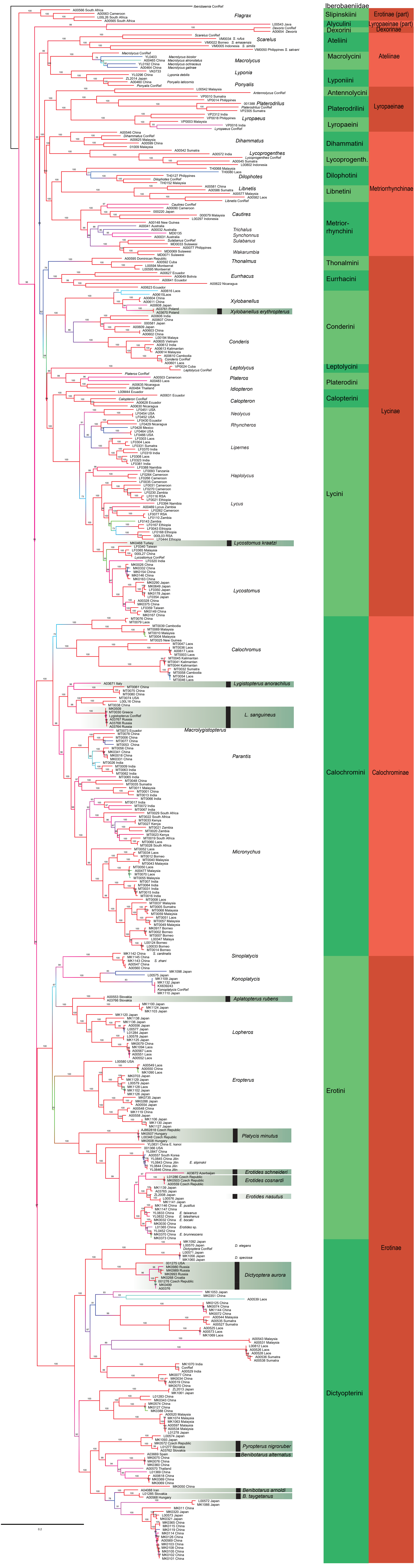


Figure S1. Molecular phylogenetic reconstruction of Lycidae relationships using maximum-likelihood.

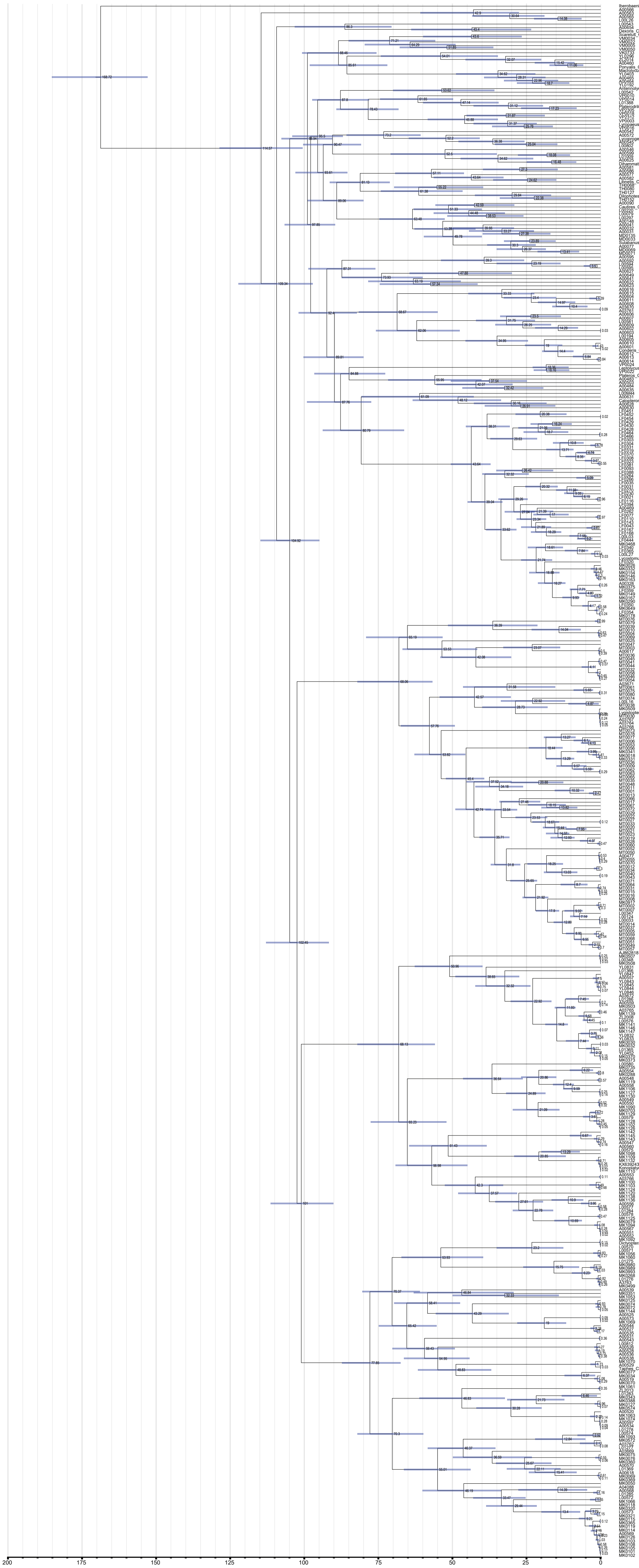


**Figure S2.** Time-calibrated, maximum clade credibility tree computed using BEAST.
